# Mechanistic implications of the interaction of the soluble substrate-binding protein with a type II ABC importer

**DOI:** 10.1101/2022.12.02.518933

**Authors:** Lingwei Zhu, Jinrang Kim, Kun Leng, John E. Ramos, Colin D. Kinz-Thompson, Nathan K. Karpowich, Ruben L. Gonzalez, John F. Hunt

**Author notes:** To whom correspondence may be addressed: JFH –, *voice*, RLG –, *voice. JK, Regeneron Pharmaceuticals Inc., Tarrytown, NY 10591, USA; KL, Medical Scientist Training Program, University of California, San Francisco, San Francisco, CA, USA; JER, Genome LLC, Hollywood, FL 33021, USA; CDK-T, Department of Chemistry, Rutgers University-Newark, Newark, NJ, 07102, USA; NKK, Structure and Protein Sciences, Janssen Pharmaceuticals, Spring House, PA, 19477, USA.

## Abstract

<u>A</u>TP-<u>B</u>inding <u>C</u>assette (ABC) Transporters employ homologous ATPase domains to drive transmembrane transport of diverse substrates ranging from small molecules to large polymers. Bacterial ABC importers require an extramembranous <u>s</u>ubstrate <u>b</u>inding <u>p</u>rotein (SBP) to deliver the transport substrate to the extracellular side of the transporter complex. Previous studies suggest significant differences in the transport mechanisms of type I *vs*. type II bacterial ABC importers, which contain unrelated transmembrane domains. We herein use ensemble fluorescence resonance energy transfer (FRET) experiments to characterize the kinetics of SBP interaction in the *E. coli* BtuCD-F complex, a canonical type II ABC importer that transports vitamin B_12_. We demonstrate that, in the absence of B_12_, BtuF (the SBP) forms a ‘locked’ (kinetically hyper-stable) complex with nanodisc-reconstituted BtuCD that can only be dissociated by ATP hydrolysis, which represents a futile reaction cycle. Notably, no type I importer has been observed to form an equivalent locked complex. We also show that either ATP or vitamin B_12_ binding substantially slows formation of the locked BtuCD-F complex, which will limit the occurrence of futile hydrolysis under physiological conditions.

Mutagenesis experiments demonstrate that efficient locking requires concerted interaction of BtuCD with residues on both sides of the B_12_ binding pocket in BtuF. Combined with the kinetic inhibition of locking by ATP binding, these observations imply that the transition state for the locking reaction involves a global alteration in the conformation of BtuCD that extends from its BtuF binding site in the periplasm to its ATP-binding sites on the opposite side of the membrane in the cytoplasm. These observations suggest that locking, which seals the extracellular B_12_ entry site of the transporter, may help push B_12_ through the transporter and directly contribute to the transport mechanism in type II ABC importers.

## 1. INTRODUCTION

ABC Transporters belong to one of the biggest protein superfamilies and can be found in organisms from all three phylogenetic kingdoms^1^. The defining characteristic of the ABC superfamily is the use of homologous homodimeric or heterodimeric ATP-binding domains, also called nucleotide-binding domains (NBDs). The conservation of these domains implies shared features in their transport mechanisms. Two ATP molecules bind and are hydrolyzed at the interface of the NBD dimer, driving conformational changes of the dimeric transmembrane domains (TMDs) that eventually lead to translocation of the substrate molecule. ABC Transporters play important roles in bacterial physiology and virulence^2^ by transporting substrates including, but not limited to, vitamins^3–5^, amino acids^6^, oligopeptides^7^, sugars^8^, and metal ions^9^. Import of iron chelation complexes by type II ABC importers makes important contributions to both bacterial fitness and pathogenesis^2^. ABC Transporters also play essential roles in eukaryotes^10–12^. In humans, upregulation of some can cause multi-drug resistance in cancer chemotherapy^13,14^, while mutations in others contribute to a variety of inherited diseases including cystic fibrosis and age-related macular degeneration^15,16^.

ABC Transporters are classified as either exporters or importers depending on the direction of movement of their transport substrates. Among the importers, they are further classified as type I, type II, and ECF (or type III) importers based on their TMD structures^17^. The diversity in TMD structures determines the wide range of substrates, many of which are dramatically different in sizes and structures, that these ABC Transporters can transport.

For type I and type II importers, substrate binding proteins (SBPs) play an essential role in substrate recognition and transport, as the transporters themselves have low or no affinity for the substrates ^18,19^. In Gram-negative bacteria, which have a periplasmic space enclosed between their inner and outer membranes, the SBPs are generally soluble proteins encoded in a separate polypeptide chain from the other components of the transporter. In Gram-positive bacteria, which only have a single membrane, the SBPs are covalently attached to a transmembrane domain in the transporter or anchored onto the membrane. The SBPs contain two asymmetrical domains (N- and C-terminal lobes) connected by a hinge region, and these two domains flank a central substrate binding site^20^ (Figure 1a). During transport, substrate-loaded SBP binds and aligns the substrate to the entrance channel of the importer. Interactions of both lobes of the SBPs with their cognate transporters are believed to play essential roles in substrate translocation. Previous studies have focused on measuring binding affinities between the importers and their cognate SBPs^4,21^. Nonetheless, little is known about how the individual lobes contribute to SBP-transporter interactions and substrate translocation.

**Figure 1.**
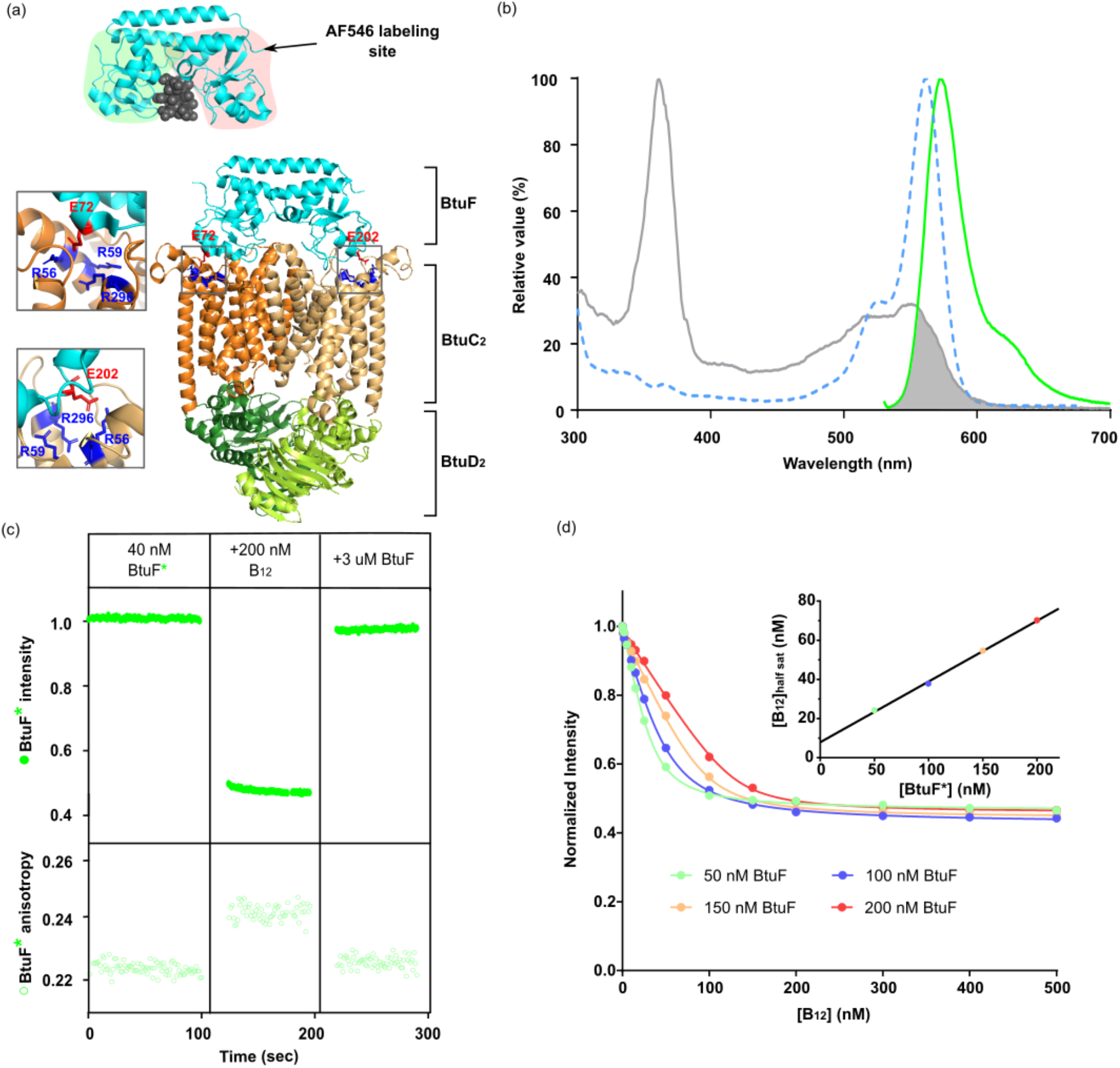
The use of fluorescence resonance energy transfer quenching to quantify vitamin B_12_ binding to AlexaFlour 546 (AF-546) labeled *E. coli* BtuF (BtuF*). **a**, The upper ribbon diagram shows a crystal structure of the BtuF-B_12_ complex (PDB: 1N4A). The N- and C-terminal lobes are indicated by green and red backgrounds, respectively. The lower ribbon diagram shows a crystal structure of the BtuCD-F complex (PDB: 2QI9). Residues E72 and E202 in BtuF are shown in red in stick representation, while residues R56, R59, and R295 in BtuC subunits are shown in blue in stick representation. **b**, The grey solid line shows absorption spectrum of B_12_ (cyanocobalamin), while the blue dash line and green solid line show excitation and emission spectra of AF546, respectively. The overlap region between absorption spectrum of B_12_ and excitation spectrum of AF546 is indicated by a grey background. **c**, Fluorescence quenching of AF546 emission from BtuF* indicates binding of B_12_, which also leads to an anisotropy increase due to the decrease of AF546 lifetime caused by FRET. BtuF*, B_12_, and unlabeled BtuF were added at the beginning of each time window at the specified concentrations. **d**, Titration of B_12_ onto various concentrations of BtuF* are shown with global fits to the quadratic binding equation. The inset plots [B_12_] at half-saturation (determined by single-curve fitting) *vs*. BtuF concentrations, a global analysis that yields a *K*_*d*_ of 7.9 ± 1.3 nM for B_12_ binding. These experiments were conducted at 25 °C in 2 mM MgCl_2_, 150 mM NaCl, 50 mM Tris•HCl, pH 7.5.

Extensive structural studies have shown that many ABC Transporters utilize an alternating access mechanism for substrate transport ^22–29^. For ABC exporters and type I importers, crystal structures show that binding of ATP leads to conformational changes that convert them from an “inward-facing” conformation to an “outward-facing” conformation, a transition that results in substrate translocation. In contrast, ATP binding to type II ABC importers does not lead to a conformational change of sufficient scale to allow substrate passage through them^19,30–32^, suggesting that transient, high-energy intermediate conformations play a critical functional role in the transport reaction. In the absence of experimental data characterizing such conformations, the mechanism of type II ABC importers remains controversial.

We have used BtuCD, an extensively characterized type II ABC importer that transports vitamin B^12 4,19,30–35^, as a model to investigate the roles of ATP binding/hydrolysis and SBP binding in substrate translocation. The TMDs of BtuCD are composed of a BtuC homodimer (Figure 1a), which contains a translocation cavity that is slightly smaller than B^1219,30^. The NBDs of BtuCD are composed of a BtuD homodimer (Figure 1a). The SBP of BtuCD is BtuF, which binds B_12_ in the periplasm. Previous studies suggest that the N- and C-terminal lobes of BtuF interact with the individual BtuC subunits through the formation of glutamic-acid/arginine salt bridges (Figure 1a). In the four available X-ray crystal structures of BtuCD or BtuCD-F^19,30–32^, the dimensions of the openings of the translocation cavity in BtuC are not large enough to allow passage of B_12_ to either the periplasmic or the cytoplasmic side of the membrane. Furthermore, it has not thus far been possible to obtain structures with B_12_ bound to either the binding site on BtuF or the cavity in BtuC_2_ within the BtuCD-F complex^4,30,33–35^. It’s transport mechanism thus remains enigmatic.

In the absence of ATP or B_12_, BtuF forms a kinetically hyper-stable complex with BtuCD that we refer to as the ‘locked’ BtuCD-F complex (Figure 1d)^4,30,35,36^. Formation of this locked complex only slightly stimulates ATP hydrolysis compared to the absence of BtuF. In contrast, the type I ABC importer for maltose, MalFGK_2_, only forms a locked complex with its SBP following ATP binding to both NBDs and prior to hydrolysis, while hydrolysis is greatly stimulated by binding of the substrate-bound SBP to the transporter^21,37^. These observations suggest that type II ABC importers employ a distinct mechanism, potentially related to the larger sizes of their substrates (*e*.*g*., 1,355 Da for B_12_) compared to most of the substrates for type I ABC importers (*e*.*g*., 342 Da for maltose).

Herein, we use fluorescence resonance energy transfer (FRET) to characterize the mechanism of SBP locking in type II ABC importers and its contribution to substrate transport. We also exploit the photophysical properties of B_12_, specifically its ability to induce FRET-based quenching of fluorophores that have an emission spectrum overlapping its absorbance spectrum^38–41^ (Figure 1b), to monitor binding of B_12_ to BtuF or the BtuCD-F complex. Our data suggest that SBP locking is directly related to the unique transport mechanism employed by these transporters. We demonstrate that high-affinity binding of both lobes of the SBP to the importer is required for locking. Our data further show that locking involves a conformational transition with high activation enthalpy in which there are coupled structural changes throughout BtuCD, extending from the SBP-binding interface in the periplasm to the ATPase active sites in the cytoplasm. We propose that the conformational transition mediating locking directly contributes to moving B_12_ through BtuCD during the transport process while simultaneously sealing the periplasmic opening of the importer to prevent back-flow.

## 2. RESULTS

### 2.1 Fluorophore-based reporter system to monitor BtuF binding to B_12_ and BtuCD

Vitamin B_12_ is a chromophore that absorbs visible light in the < 600 nm wavelength region but has no detectable fluorescence emission. It can therefore serve as a FRET acceptor for a donor with fluorescence emission in this wavelength range, although the effect of this energy transfer will only be observed via quenching of the FRET donor fluorescence^38^ (*i*.*e*., a reduction in its fluorescence emission intensity without any corresponding fluorescence emission from the B_12_ acceptor). Overlap (Figure 1b) between the B_12_ absorption spectrum and the emission spectrum of the commonly used fluorophore Alexa Fluor 546 (AF546) thus enables FRET-induced quenching to be exploited to monitor the distance between B_12_ and AF546-labeled BtuF, which we call BtuF* (Figure 1c-d). Note that emission intensity is monitored exclusively from the donor fluorophore in these ‘single-color’ FRET-quenching measurements in which only BtuF* is labeled, in contrast to the common ‘two-color’ FRET experiments in which an increase in acceptor fluorescence is monitored in parallel with a decrease in donor fluorescence. Such two-color FRET experiments are also reported below in which BtuCD is labeled with AF647 (BtuCD*) to characterize its interaction with AF546-labeled BtuF* in the absence of B_12_ ; the absorption spectrum of AF647 overlaps the emission spectrum of AF546, but, unlike B_12_, AF647 emits fluorescence upon resonance energy transfer from the AF546 donor, enabling the BtuF*-BtuCD* interaction to be monitored using standard two-color FRET methods.

Our single-color FRET quenching experiments show that binding of B_12_ to BtuF* (Figure 1d) leads to ∼55% quenching of AF546 emission intensity (Figure 1b-c). It simultaneously produces an ∼0.02 increase in the fluorescence anisotropy of BtuF* (first two time segments in Figure 1c) because FRET-induced quenching decreases the fluorescence lifetime of AF546. The slower rotational diffusion of the BtuCD-F* complex compared to free BtuF* produces a larger change in BtuF* anisotropy upon binding to BtuCD, which enables the BtuF* fluorescence signal to be used to monitor both B_12_ binding and BtuCD binding simultaneously, as explained further below.

Addition of unlabeled BtuF to a solution with BtuF* saturated with B_12_ leads within 20 sec to complete reversal of both the fluorescence and anisotropy changes observed upon B_12_ binding to BtuCD (final time segment in Figure 1c), indicating that B_12_ exchanges rapidly between BtuF-bound and BtuF-free states. Global fitting of binding experiments conducted at four different BtuF* concentrations indicates that the equilibrium dissociation constant (K_d_) for binding of B_12_ to BtuF* is 7.9 nM (Figure 1d), matching published estimates and isothermal titration calorimetry (ITC) data on unlabeled BtuF (Figure S1). The ITC data show that the detergent lauryldimethylamine oxide (LDAO) blocks the binding of B_12_ to BtuF (which is one reasons that we conducted the mechanistic experiments described below with BtuCD reconstituted into nanodiscs rather than solubilized in detergent).

### 2.2 Formation in the absence of ATP of a locked BtuCD-F* complex that cannot bind B_12_

Previous studies have shown that, in the absence of ATP or B_12,_ *apo*-BtuF binds directly to BtuCD with extremely high affinity, resulting in formation of a ‘locked’ (kinetically hyper-stable) BtuCD-F complex that was hypothesized to be destabilized by either B_12_ or ATP interaction^4,30,35,36^. However, the exact molecular mechanism driving dissociation of the locked complex remains unclear, including whether its dissociation is driven by ATP binding *vs*. hydrolysis. We exploited the fluorescence anisotropy of BtuF* to characterize the mechanistic features of the locking reaction in greater detail (Figures 2-5). Because of the large reduction in the rotational diffusion coefficient of BtuF* upon formation of the BtuCD-F* complex, the anisotropy of the BtuF* fluorescence emission increases significantly upon complex formation, while its intensity remains unchanged (Figure 2a). Because BtuCD and B_12_ increase BtuF* anisotropy via different physical mechanisms, the two binding reactions make independent contributions to anisotropy. Furthermore, because B_12_ binding quenches emission intensity (Figure 1c) while BtuCD binding has no influence on it (Figure 2a), monitoring the fluorescence emission intensity of BtuF*in parallel with its anisotropy enables BtuCD and B_12_ binding events to be distinguished.

**Figure 2.**
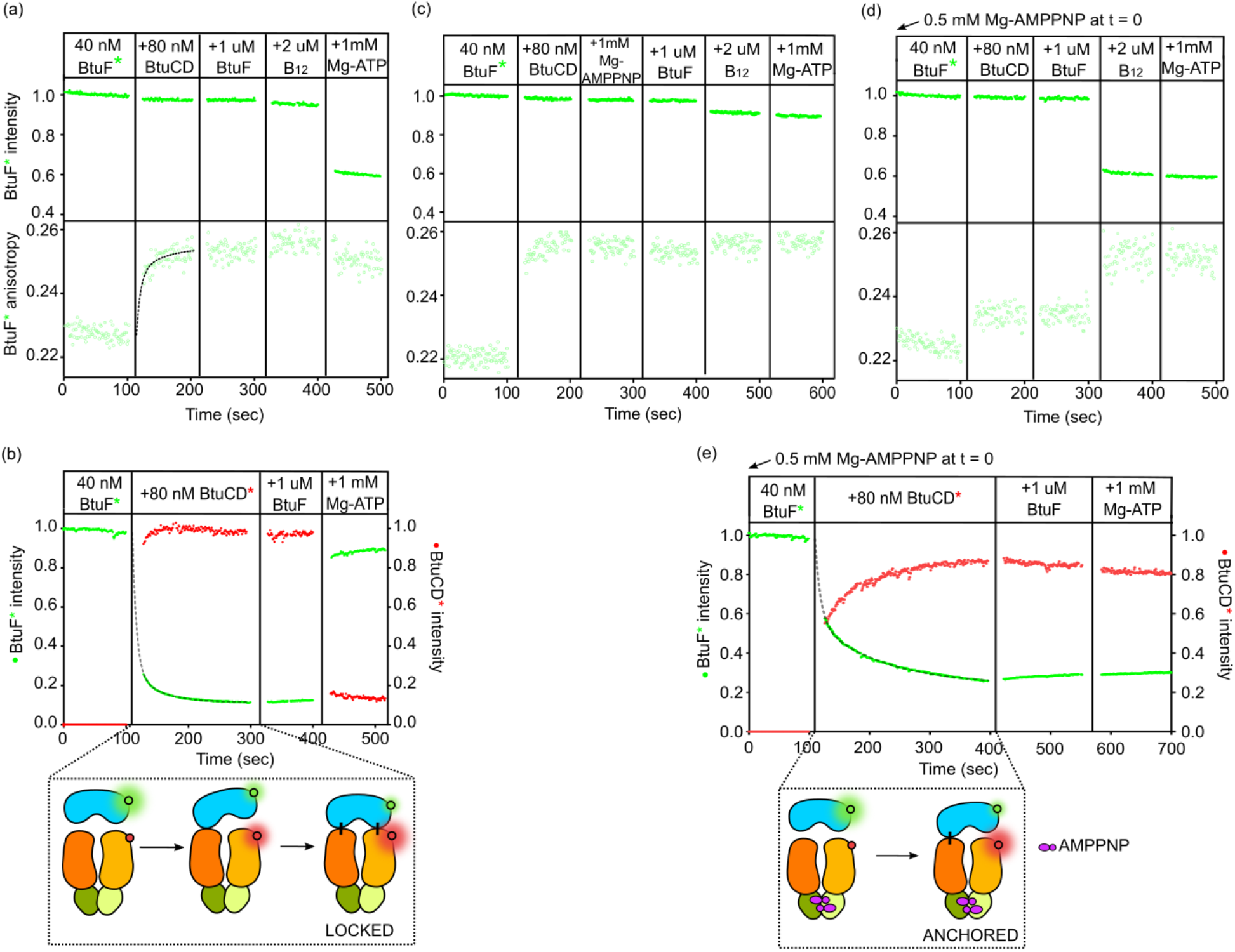
BtuF* forms a locked complex with BtuCD in the absence of ATP that prevents B_12_ binding, while it forms an anchored complex with BtuCD•AMPPNP that allows B_12_ binding. **a**,**c**,**d**, Experiments are shown using simultaneous fluorescence and anisotropy measurements to monitor interaction between BtuF* and unlabeled BtuCD. Formation of BtuCD-F* complex is indicated by the increase in anisotropy upon the addition of BtuCD. The anisotropy is fitted with a double exponential function using parameters obtained from the curve-fit in panel **b**. No change of anisotropy was observed upon the addition of excess unlabeled BtuF to the locked BtuCD-F* complex. Minimal quenching of fluorescence emission was observed upon the addition of B_12_. Panel **c** shows an experiment similar to panel **a**, but with the addition of AMPPNP prior to the addition of unlabeled BtuF. Panel **d** shows an experiment similar to panel **a**, but with the inclusion of 0.5 mM AMPPNP at t = 0 sec. The anchored BtuCD•AMPPNP-BtuF* complex was formed upon the addition of BtuCD•AMPPNP to BtuF*, with smaller change of anisotropy compared to the experiment shown in panel **a**. Addition of excess unlabeled BtuF showed no change of anisotropy, which indicates formation of stable complex. Addition of B_12_ leads to significant quenching, indicating binding of B_12_ to the anchored BtuCD-F* complex. **b**,**e**, Two-color ensemble binding experiment are shown using AF546 labeled BtuF* (green) and AF647 labeled BtuCD* (red) to monitor their interaction in the absence of B_12_. Binding of BtuF* to BtuCD* is indicated by the increase of red emission and the decrease of green emission. Change of green emission upon the addition of BtuCD* was curve-fit by a double exponential function, yielding k_fast_ = 0.158 ± 0.002 sec^-1^ and k_slow_ = 0.025 ± 0.0004 sec^-1^ (corresponding to time constants of 6.3 and 40 sec, respectively). The fast rate constant represents the initial single-lobe association between BtuF* and BtuCD, while the slow rate constant represents the locking of BtuCD-F* complex, as illustrated in the schematic. Panel **e** shows an experiment similar to panel **b**, but with the inclusion of 0.5 mM AMPPNP at t = 0 sec. Double exponential curve-fitting of these data yields k_fast_ = 0.108 ± 0.002 sec^-1^ and k_slow_ = 0.009 ± 0.0001 sec^-1^ (corresponding to time constants of 9.3 and 111 sec, respectively). The experiments were conducted at 25 °C in the same buffer used in Figure 1.

Single-color ensemble experiments show an ∼0.025 increase in the anisotropy of BtuF* fluorescence emission but no change in its intensity upon addition of 80 nM unlabeled BtuCD reconstituted into MSP1E3D1^42^ nanodiscs with *E. coli* polar lipids (first two time segments in Figure 2a). These data are consistent with nearly stoichiometric of binding of BtuF* to BtuCD with a time constant of ∼40 sec. Addition of a 25-fold excess unlabeled BtuF to the resulting BtuCD-F* complex after 100 seconds produces no significant change in anisotropy (third time segment in Figure 2a), indicating that, on this time scale, the BtuCD-bound BtuF* does exchange with the BtuF free in solution. Therefore, formation of the BtuCD-F* complex is effectively kinetically irreversible; this observation led us to designate this complex as ‘locked’.

Subsequent addition of B_12_ to the locked complex results in no change in either the anisotropy or the intensity of the fluorescence emission from BtuF* (fourth time segment in Figure 2a) until ATP is added (final time segment in Figure 2a), showing that formation of the locked BtuCD-F* complex prevents B_12_ from binding until the addition of ATP. This conclusion is consistent with the X-ray crystal structure of the BtuCD-F complex in which the B_12_ binding site in BtuF is partially occupied by periplasmic residues from BtuC_2_^31^. Note that addition of ATP to the locked BtuCD-F complex rapidly produces strong quenching of BtuF* fluorescence (final time segment in Figure 2a), indicating that ATP binding or hydrolysis by BtuCD restored BtuF* to a state in which it can bind B_12_. This result suggests that addition of ATP dissociates the locked complex, which exposes the B_12_ binding site on BtuF* when it is released from the complex. The quenching of BtuF* fluorescence intensity by B_12_ in Figure 2a is accompanied by only a small decrease in anisotropy because the reduction in fluorescence lifetime due to quenching is largely offset by the increase in the rotational diffusion rate of BtuF* when it is released from BtuCD.

A two-color ensemble FRET experiment using AF546-labeled BtuF* as the donor and AF647-labeled BtuCD* as the acceptor (as described above) confirms that the interaction of BtuF with BtuCD is effectively irreversible in the absence of ATP or B_12_ (first three time segments in Figure 2b), and this experiment also corroborates the conclusion that the locked complex is fully dissociated by ATP (final time segments in Figure 2b). Curve fitting analysis of the reduction in BtuF* intensity upon the addition of BtuCD* in the double-emission FRET experiment indicates that the association reaction (second time segment in Figure 2b) is composed of a fast phase with a time constant of 6.3 sec followed by a slow phase with a time constant of 40 sec, with the latter matching that of the anisotropy change in the single-emission FRET-quenching experiment described above (Figure 2a). The fast phase seems likely to reflect initial docking of BtuF* to BtuCD* in a flexibly-bound state, probably via one of its two lobes, whereas the slow phase seems likely to represent the locking reaction that involves both of the lobes, based on additional data presented below.

Another single-color FRET-quenching experiment shows that adding AMPPNP, a nonhydrolyzable analog of ATP, to the preformed locked BtuCD-F* complex (first two time segments in Figure 2c) does not produce a detectable change in anisotropy (third time segment in Figure 2c), indicating that binding of the ATP analog does not induce significant dissociation of the locked complex. Taking this result together with the observation that addition of ATP does induce dissociation of the locked complex (final time segment in Figure 2a) supports ATP hydrolysis, instead of ATP binding, driving complex dissociation. Therefore, ATP hydrolysis that is uncoupled from transport is required for dissociation of the locked BtuCD-F* complex. Type I ABC importers do not exhibit any equivalent locking phenomenon ^8,21,37,43^, reinforcing the inference that Type II ABC importers employ a significantly different transport mechanism.

While the mechanistic function of SBP locking to type II ABC importers has been the subject of speculation in previous studies^4,30,35,36^, it has not been clearly explained. The ATP hydrolysis that we demonstrate here to be required for dissociation of the locked SBP from the transporter would be thermodynamically futile if the complex does not play a significant role in the progression or regulation of the B_12_ transport reaction.

### 2.3 Formation in the presence of AMPPNP of an anchored BtuCD-F* complex that still binds B_12_

In *E. coli*, the physiological concentration of ATP is typically in the millimolar range^44–47^. Given that the previously reported Michaelis constant, K_m_, for ATP hydrolysis by BtuCD is ∼20 µM^3^, BtuCD should be saturated by ATP *in vivo* when binding BtuF. Therefore, the interaction between BtuF and BtuCD•AMPPNP represents a model for the initial stage of the transport reaction. In single-color FRET-quenching experiments, addition of the preformed BtuCD•AMPPNP complex to BtuF* (first two time segments in Figure 2d) produces a significantly smaller (∼0.01 vs. ∼0.025) change in anisotropy than when BtuCD is added to BtuF* in the absence of any nucleotide (first two time segments in Figure 2a). This result indicates that the interaction of BtuF* with BtuCD•AMPPNP is significantly altered in some way relative to its interaction with nucleotide-free BtuCD, an inference confirmed by the B_12_-binding properties established below for this complex compared to the complex in the absence of nucleotide. Subsequent addition of a 25-fold excess of unlabeled BtuF does not affect the anisotropy of BtuF*, indicating that the complex formed in the presence of AMPPNP is also effectively kinetically irreversible because it shows negligible dissociation during the 100 sec observation period (third time segment in Figure 2d). We call this kinetically irreversible complex ‘anchored’ to distinguish it from the structurally and mechanistically distinct locked BtuCD-F* complex formed in the absence of nucleotide.

In contrast to the result observed for the BtuCD-F* locked complex (fourth time segment in Figure 2a), addition of B_12_ to the BtuCD•AMPPNP-BtuF* anchored complex results in ∼40% quenching of the fluorescence of BtuF* (fourth time segment in Figure 2d), indicating that the B_12_ binding site in BtuF* is mostly accessible in the anchored complex. This difference indicates a substantial difference in conformation in the complex formed in the presence of AMPPNP, which, as noted above, is likely to be a model of the initial functional complex formed during the B_12_ transport reaction given the high concentration of ATP present *in vivo*. Finally, addition of 1 mM ATP to the anchored BtuCD•AMPPNP-ButF* complex does not result in a change in either BtuF* intensity or anisotropy (final time segment in Figure 2d), indicating that BtuF* does not dissociate under these conditions, in contrast to the effect of ATP addition to the locked BtuCD-F* complex in the absence of nucleotide. Therefore, AMPPNP is unable to exchange with free nucleotide when bound in the BtuCD•AMPPNP-BtuF* anchored complex, meaning AMPPNP is also effectively irreversibly bound in this complex.

In a two-color ensemble FRET experiment, addition of the preformed BtuCD*•AMPPNP complex to BtuF* (first two time segments in Figure 2e) initially results in only half of the BtuF* intensity increase due to FRET relative to that observed in the equivalent experiment performed in the absence of nucleotide (first two time segments in Figure 2b). This initial BtuF* intensity increase occurs rapidly, with a time constant of 9.3 sec, and is followed by a much slower increase in FRET with a time constant of 111 sec. We conclude that this slow phase of the BtuF* intensity increase in the two-color FRET experiment represents a conformational relaxation of some kind within the anchored BtuCD*•AMPPNP-BtuF* complex that does not affect the mobility of BtuF* because the equivalent single-color FRET-quenching experiment shows that the change in BtuF* anisotropy is completed on a substantially faster time scale (first two time segments in Figure 2a). Subsequent addition of a 25-fold excess of unlabeled BtuF (third time segment in Figure 2e) and then 1 mM ATP (final time segment in Figure 2e) to the anchored BtuCD*•AMPPNP-BtuF* complex both produce no change in BtuF* intensity, confirming the inference from the equivalent single-color emission FRET-quenching experiment (final two time segments in Figure 2d) that BtuF* and AMPPNP are both effectively bound irreversibly in the anchored BtuCD*-BtuF*-AMPPNP complex.

The accessibility of the B_12_ binding site in the anchored BtuCD•AMPPNP-BtuF* complex (fourth time segment in Figure 2d) suggests BtuCD is interacting with BtuF* on only one of the two lobes surrounding its central B_12_ binding site (Figure 1d) because interaction with both lobes would sterically block that site. Previous studies of type I and type II ABC importers have hypothesized the formation of such an SBP-transporter complex involving single-lobe binding of the SBP^35,39,48,49^ without direct experimental support. As noted above, the change of anisotropy upon the addition of BtuCD to BtuF* in our single-color FRET-quenching experiments is substantially smaller in the anchored BtuCD•AMPPNP-BtuF* complex compared to the locked BtuCD-F* complex formed in the absence of nucleotide (compare Figure 2d *vs*. Figure 2a), indicating higher mobility of BtuF* within the anchored complex, despite its kinetically stable binding. We therefore propose that only one of the two lobes of BtuF* is tightly associated with BtuCD in this complex, allowing BtuF* to flexibly sample conformations that expose its B_12_ binding site to solution. We report below the results of mutagenesis experiments strongly supporting this hypothesis.

### 2.4 Binding of B_12_ to BtuF* blocks formation of the locked BtuCD-F* complex

We then used single-color FRET-quenching experiments to characterize how binding of B_12_ to BtuF* influences the locking reaction (Figures 3 & S2). Adding BtuCD to the preformed BtuF*-B_12_ complex leads to gradual dequenching of AF546 intensity over the course of several hundred seconds, with the rate of dequenching progressively slowing as the concentration of free B_12_ is increased (third time segment in Figure 3a). Quantitative analysis of the rate of dequenching as a function of B_12_ concentration (inset in Figure 3a) suggests this dequenching process represents formation of the locked BtuCD-F* complex exclusively during transient release of B_12_ from BtuF*. We use the term “trapping” to describe this kinetic process in which the effectively irreversible locking reaction occurs during transient release of B_12_ from BtuF* (as schematized at the bottom of Figure 3a), leading to a slow unidirectional progression of BtuF* into the B_12_-free unquenched state as it is trapped in the locked BtuCD-F* complex. (Note that the terms trapped/trapping and locked/locking are used with these different meanings throughout this manuscript.)

**Figure 3.**
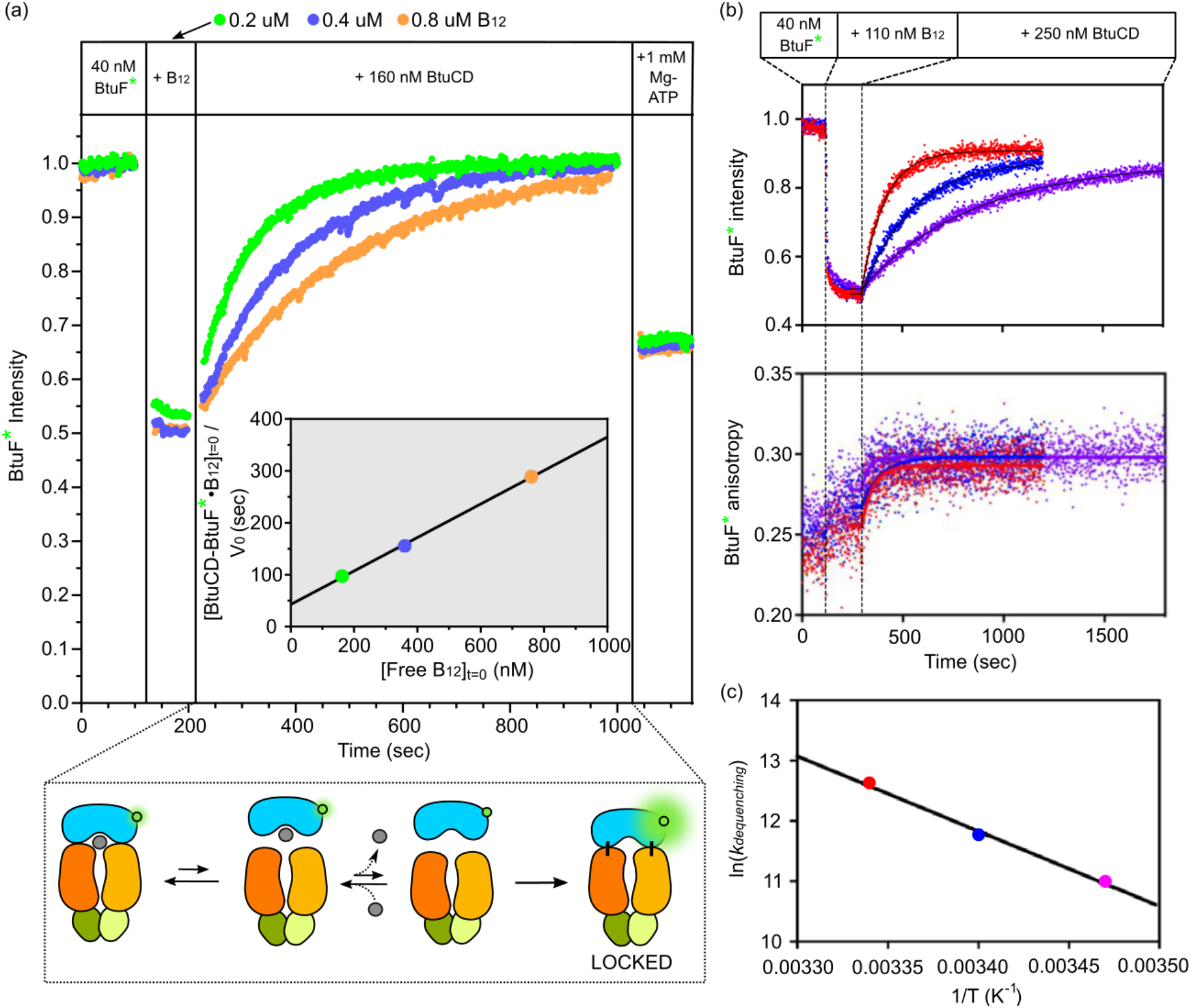
B_12_ binding to BtuF* prevents locking to BtuCD. **a**, Experiments adding 0.2 uM (green), 0.4 uM (blue), or 0.8 uM (orange) B_12_ to BtuF* prior to the additions of constant concentrations of unlabeled BtuCD and ATP. The schematic shows *apo*-BtuF* being captured by BtuCD and forming a locked BtuCD-F* complex that prevents B_12_ rebinding, which is manifested as dequenching of AF546 fluorescence. The experiment was conducted at 25° C in the same buffer used in Figure 1. The inset shows a plot of [BtuCD-F*-B_12_]_t=0_/V_0_ vs. [free B_12_]_t=0_, which should be linear for the trapping mechanism illustrated in the schematic based on the equation describing this process derived in the **Methods** section. **b**, Three experiments are shown in which BtuF*, B_12_, and BtuCD were added at indicated time points to samples at 15° C (purple), 20° C (blue), or 25 ° C (red) in the same buffer used in Figure 1. **c**, An Arrhenius plot is shown derived from single-exponential curve-fitting of the data shown in panel **b**. The estimated activation enthalpy for BtuCD-F locking is 103 ± 6 kJ/mol.

A simple steady-state model for the kinetics of the trapping process predicts the concentration-normalized initial rate of the reaction should exhibit a linear dependence on free B_12_ concentration (see Methods), and the observed kinetics of the trapping process are consistent with this prediction (inset in Figure 3a). This analysis indicates B_12_ is released from the reversibly associated BtuCD-BtuF* complex at a rate 0.01 sec^-1^, approximately one order-of-magnitude slower than the release rate from BtuF* not bound to BtuCD as measured using single-molecule, single-color FRET-quenching measurements^50^. A reduction in release rate is consistent with a restriction in the movement of B_12_ bound to BtuF* when its B_12_-binding site is closely apposed to the interface of BtuCD in the reversibly associated complex.

Repeating the same trapping experiment at different temperatures (Figure 3b) enables the Arrhenius equation to be used to estimate an activation enthalpy of ∼103 kJ/mol (Figure 3c) for the locking reaction (*i*.*e*., the irreversible step in the trapping process). An activation enthalpy of this magnitude, which requires large changes in interatomic interactions, is consistent with a substantial difference in protein conformation in the transition state for the reaction compared to the pre-reaction complex.

Locking of BtuF* to BtuCD exclusively upon release of B_12_ is further supported by a series of competition experiments in which a 120-fold excess of unlabeled BtuF is added at different times after initiating the trapping process (Figure S2a). This addition stops locking of BtuF*, due to competition for binding to BtuCD, and simultaneously removes B_12_ from BtuF*, due to competition for binding of released B_12_. Quantitative analysis of the intensity and anisotropy data from these experiments shows that the proportion of the BtuF* population in the unlocked state during the trapping process is equal to the proportion with B_12_ bound (Figure S2b), again consistent with locking being tightly coupled to release of B_12_ from BtuF*.

Repetition of the same single-color FRET-quenching experiment in the presence of AMPPNP demonstrates that AMPPNP stops the trapping process (Figure S3), consistent with the data above showing binding of this nonhydrolyzable ATP analog to BtuCD blocks formation of the locked BtuCD-F* complex (Figure 2). The stable BtuCD•AMPPNP-BtuF*•B_12_ complex formed under these conditions in Figure S3 provides a model for the fully formed functional transporter complex with ATP bound but prior to ATP hydrolysis. Notably, the BtuF* intensity level in this complex indicates B_12_ remains bound to its binding site on BtuF* with little movement compared to its location prior to BtuF*•B_12_ binding to BtuCD, strongly supporting ATP hydrolysis being needed for B_12_ movement into the transport channel in BtuCD.

### 2.5 Characterization of single-lobe interactions between BtuF* and BtuCD

Based on the results reported above, we hypothesize that locking occurs via a rapid initial binding of a single lobe of BtuF* to BtuCD followed by a slower binding at the second lobe because that interaction requires a substantial conformational change in BtuCD that is reflected in the high activation enthalpy of the locking reaction (Figure 3c). This hypothesis is supported by the two-phase kinetics observed in the two-color FRET experiment characterizing the BtuCD-F* locking reaction (Figure 2b) together with the observed properties of the anchored BtuCD•AMPPNP-BtuF* complex (Figure 2d-e). To test the hypothesis, we produced BtuF* variants harboring mutations in each of its two lobes that block BtuCD interaction at that site.

Crystal structures of the BtuCD-F complex^19,31^ show one glutamic acid on each of the two lobes of BtuF (E72 in N-terminal lobe and E202 in C-terminal lobe) inserted into an arginine “pocket” in BtuC subunits that includes R56, R59, and R295 (Figure 1d). Sequence alignment showed that these glutamic acid and arginine residues in BtuF and BtuC are all highly conserved in siderophore and cobalamin transporters in different organisms^36^. *In vivo* transport experiments have demonstrated mutation of E72 in BtuF leads to almost complete loss of transport activity, whereas mutation of E202 leads to only partial reduction of transport activity^51^, which suggests asymmetric contribution of the two lobes of BtuF to BtuCD binding. We therefore conducted ensemble fluorescence experiments equivalent to those reported above but with BtuF* variants harboring either mutations E72A or E202A mutation in order to characterize this asymmetry in more detail (Figures 4, 5, & S4).

**Figure 4.**
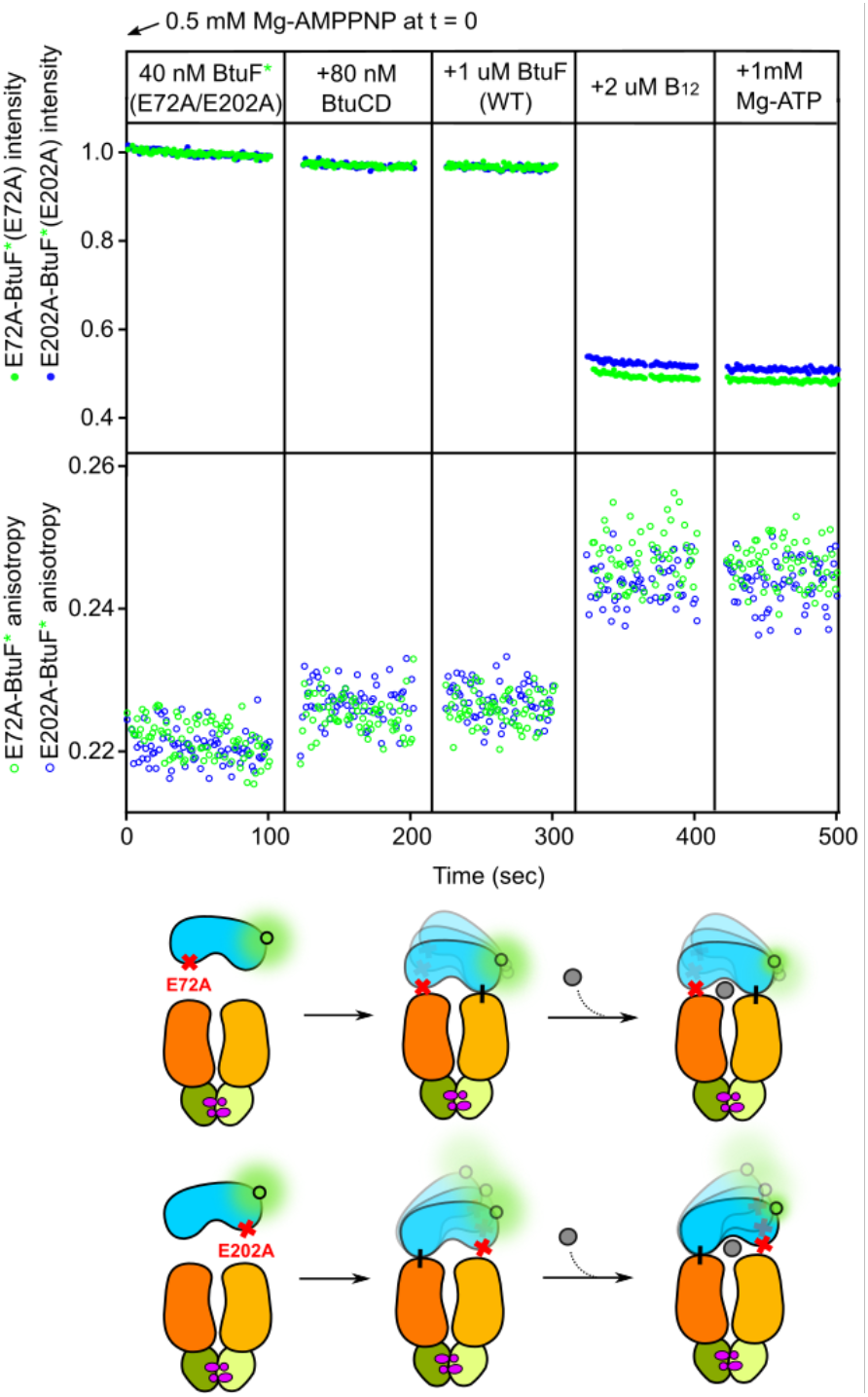
Mutagenesis of the two BtuCD interaction sites on BtuF* demonstrates that binding at either site stably anchors BtuF* to BtuCD in a conformation in which vitamin B_12_ efficiently binds to BtuF*. Experiments similar to Figure 2d, but using AF546-labeled BtuF* harboring the E72A (green) or E202A (blue) mutation. The schematic below the graph illustrates formation of a BtuCD-F* complex anchored at either lobe, but with BtuF in a flexible conformations that allows B_12_ binding. The experiments were conducted at 25° C in the same buffer used in Figure 1 but with inclusion of 0.5 mM AMPPNP.

**Figure 5.**
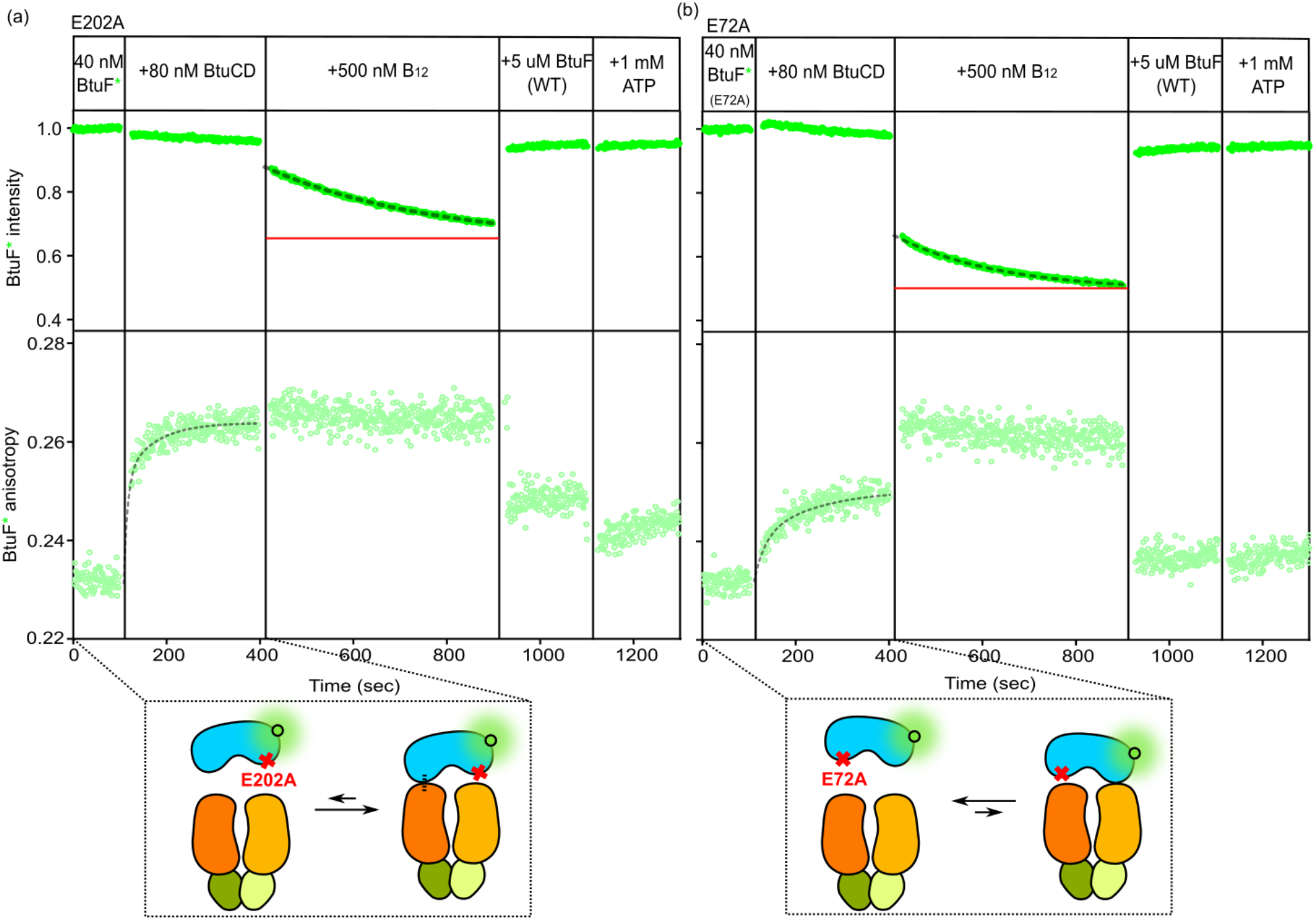
The two lobes of BtuF* make different energetic contributions to locking to BtuCD. **a**, An experiment is shown in which E202A-BtuF* and AF647-labeled BtuCD (BtuCD*) are mixed first prior to additions of B_12_, unlabeled WT-BtuF, and finally ATP. The change in anisotropy upon the addition of BtuCD was curve-fit by a double exponential function, yielding k_fast_ = 0.124 ± 0.036 sec^-1^ and k_slow_ = 0.016 ± 0.003 sec^-1^ (corresponding to time constants of 8.1 and 28 sec, respectively). The change of fluorescence intensity upon the addition of B_12_ was curve-fit by a single exponential function, yielding k = 0.0028 ± 0.0001 sec^-1^ (corresponding to a time constant of 360 sec) and plateau intensity of 0.641 ± 0.002. **b**, An experiment is shown that is equivalent to that in panel **a** but using E72A-BtuF*. The change in anisotropy upon the addition of BtuCD was curve-fit by double exponential function, yielding k_fast_ = 0.044 ± 0.019 sec^-1^ and k_slow_ = 0.008 ± 0.005 sec^-1^ (corresponding to time constants of 23 and 125 sec, respectively). The change of fluorescence intensity upon the addition of B_12_ was curve-fit by single exponential function, yielding k = 0.0045 ± 0.0001 sec^-1^ (corresponding to a time constant of 222 sec) and plateau intensity of 0.511 ± 0.001. Red horizontal lines indicate the plateau levels of the intensities from each experiment. The relatively constant BtuF* anisotropy levels in the third time segment in each panel masks offsetting changes in fluorescence lifetime due to quenching by B_12_ and rotational diffusion constant due to alterations in the interaction with BtuCD. The higher plateau level of the intensity observed upon the addition of B_12_ in the experiment using E202A-BtuF* compared to the experiment using E72A-BtuF* indicates that E202A-BtuF* that forms complex with BtuCD that can temporarily convert to conformations that block B_12_ binding.

In single-color FRET-quenching experiments conducted in the presence of AMPPNP (Figure 4), the BtuF* variants harboring either the E72A or E202A mutation behave almost identically to one another and extremely similarly to the wild-type (WT) protein except for showing slightly lower anisotropy increases in the anchored BtuCD•AMPPNP-BtuF* complexes (compare Figure 4 to Figure 2d). Following binding to BtuCD•AMPPNP, no change in anisotropy is observed for either mutant BtuF* variant upon addition of a 25-fold excess of unlabeled BtuF (third time segment in Figure 4), showing they are effectively irreversibly bound to BtuCD just like the WT protein. They also both bind B_12_ in this state (fourth time segment in Figure 4).

Therefore, both E72A-BtuF* and E202A-BtuF* form an anchored complex with BtuCD•AMPPNP with functional properties equivalent to WT BtuF* (Figure 2d). These results support either lobe of BtuF* being able to mediate tight, single-lobe anchoring to BtuCD in the initial stage of either the locking reaction or the B_12_ transport reaction. The slightly lower anisotropy levels of the mutant proteins in the anchored BtuCD•AMPPNP-BtuF* complexes suggest they have somewhat higher mobility in this state than the WT protein due to a weaker reversible interaction of the mutant lobe with BtuCD•AMPPNP when the WT lobe is irreversibly bound.

In contrast to these observations on the anchored complex, experiments directly characterizing the locking reaction demonstrate that the two lobes of BtuF* have a significantly different influence on that reaction (Figures 5 & S4). Single-color FRET-quenching experiments characterizing BtuF* locking to BtuCD in the absence of nucleotide demonstrate that E202A-BtuF* forms a complex with BtuCD slightly more slowly but with similar anisotropy properties to the locked BtuCD-F* complex formed by WT-BtuF* (first two time segments in Figure 5a).

Addition of B_12_ to this complex (third time segment in Figure 5a) and two-color FRET experiments on the same mutant protein (Figure S4a) both support ∼80% of E202A-BtuF* having its B_12_ binding site in an inaccessible state in a stable complex with BtuCD that dissociates with a time constant of ∼540 sec. Therefore, the E202A mutation in the C-terminal lobe of BtuF* significantly reduces both the efficiency of formation and the stability of the locked BtuCD-BtuF* complex formed in the absence of nucleotide.

The E72A mutation in the N-terminal lobe has qualitatively equivalent but stronger effects. Single-color FRET quenching experiments show that E72A-BtuF* forms a complex with BtuCD significantly more slowly and with only approximately half the limiting anisotropy as WT-BtuF* (third time segment in Figure 5a), while addition of B_12_ to this complex (third time segment in Figure 5b) and two-color FRET experiments on this mutant protein (Figure S4b) both support ∼20% of E72A-BtuF* being stably bound to BtuCD and released with a time constant of ∼220 sec. Therefore, mutations in either lobe of BtuF* significantly destabilize the locked BtuCD-F* complex, but the E72A mutation in the N-terminal lobe has a stronger destabilizing effect than the E202A mutation in the C-terminal lobe. These observations suggest that the N-terminal lobe both binds BtuCD faster and contributes more energy to stabilization of the locked complex than the C-terminal lobe. Therefore, the locking reaction is likely to be initiated by interaction of the N-terminal lobe of BtuF* with BtuCD followed by slower interaction of the C-terminal lobe of BtuF* to seal the complex and prevent B_12_ binding.

## 3 Discussion

We employed ensemble FRET experiments to study how interactions between BtuF and BtuCD are modulated by B_12_ or AMPPNP binding. We exploited a photophysical property of B_12_, its ability to induce FRET-quenching of AF546, to monitor its binding to AF546-labeled BtuF (BtuF*) in isolation or bound to BtuCD. Our results show that, with a non-hydrolyzable ATP analog (AMPPNP) bound to BtuCD, either lobe of BtuF can be tightly anchored to BtuCD-AMPPNP but with BtuF in a mobile conformation that still allows binding of B_12_ (Figure 6a).

**Figure 6.**
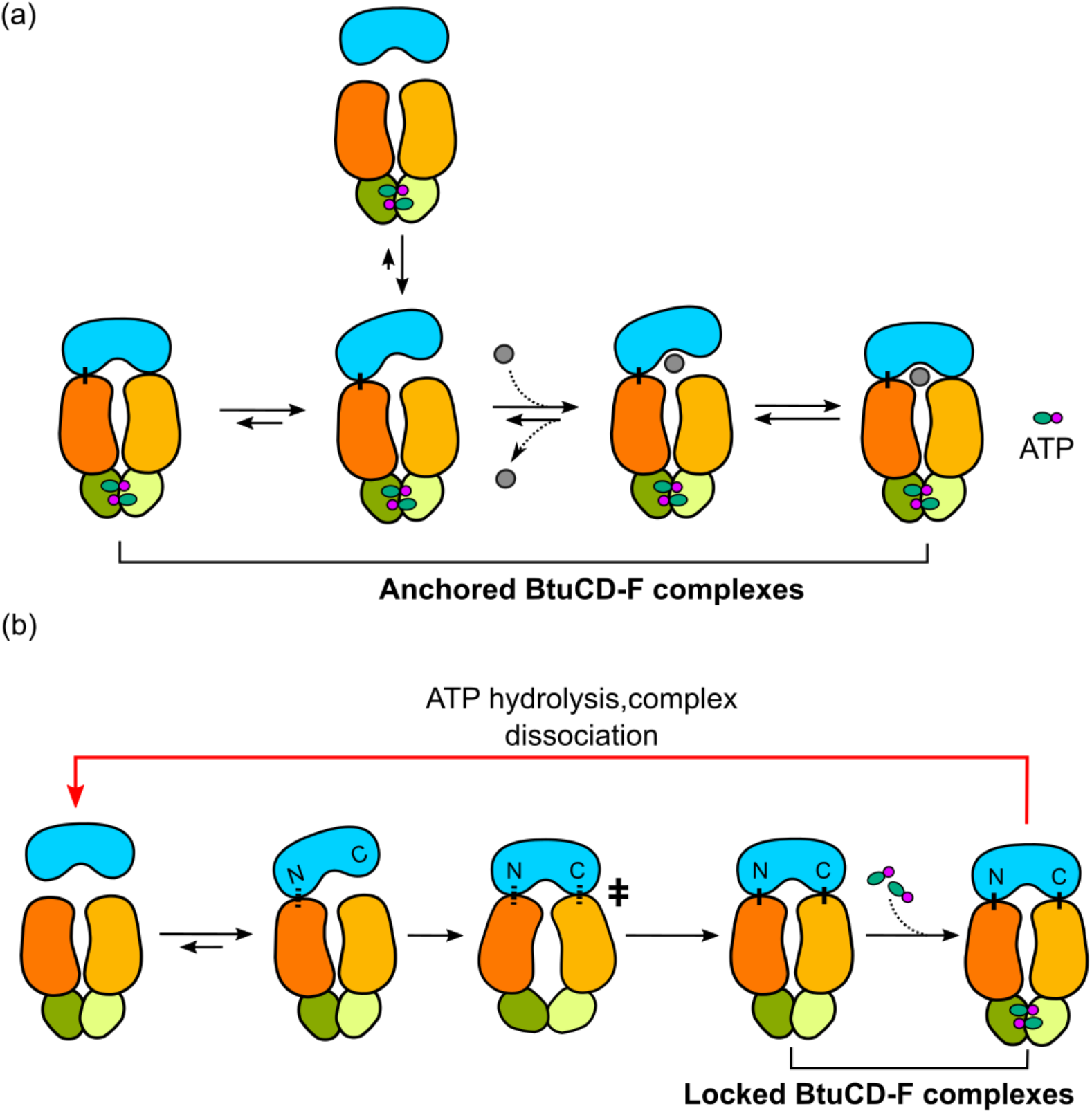
Mechanistic schematics. **a**, Schematic illustrating interactions between BtuF and BtuCD in the when ATP is bound to BtuCD. BtuF can be tightly anchored onto BtuCD with either lobe in flexible conformations that allow B_12_ binding to the complex. **b**, Schematic showing interactions between BtuF and BtuCD in the absence of ATP binding. The C-terminal lobe of BtuF (E72) first interacts with one of the BtuC subunits, which is followed by global conformational changes of BtuCD and formation of two stable salt bridges between the two lobes of BtuF and the BtuC subunits, leading to formation of the locked BtuCD-F complex that can only be dissociated by ATP hydrolysis.

Formation of such a single lobe-anchored SBP-transporter complex that can still bind to substrates has been proposed in prior studies of BtuCD^39,48^ and MalFGK_2_^49^, but not experimentally demonstrated until now. Interestingly, ABC importers with one SBP lobe fused to the transporter or anchored in the lipid bilayer have been found in gram-positive bacteria and archaea ^20,52,53^. Evolution has not produced equivalent covalent tethering of SBPs in gram-negative bacteria, presumably because they are sequestered in the periplasmic space between the inner and outer membranes where they can bind substrates with high affinity. However, the results reported here show that very strong SBP-transporter interactions in the absence of substrate have been maintained, at least in type II importers, suggesting this interaction is likely to play an important role in the transport mechanism.

We also show that, in the absence of ATP, the N-terminal lobe of BtuF containing residue E72 interacts more strongly with BtuCD than the C-terminal lobe containing residue E202. Both lobes, however, are required for the formation of the locked BtuCD-F complex that completely prevents B_12_ binding and BtuF dissociation from BtuCD. We therefore propose that, initially, the N-terminal lobe of BtuF interacts with one BtuC subunit, an event that is followed by interaction of the C-terminal lobe with the other BtuC subunit, leading to formation of the locked complex. Importantly, the locked complex cannot dissociate until ATP is hydrolyzed (Figures 2a-c & 6b). This uneven contribution of N- and C-terminal lobes of BtuF to locking is consistent with the asymmetry observed in the crystal structure of the BtuCD-F complex and with a previous functional study^51^ in which E202A-BtuF mutant showed an ∼2.5-fold reduction in transport activity while E72A-BtuF (E72A) mutant showed a complete loss of activity.

We additionally demonstrate that binding of B_12_ to BtuF (Figures 3 & S2) or binding of the non-hydrolyzable ATP analog AMPPNP to BtuCD (Figures 2d, 2e, & S3) both effectively block formation of the locked BtuCD-F complex. Crystal structures show that binding of AMPPNP to BtuCD leads to slight closure of the transmembrane cavity at the interface of the symmetrical BtuC subunits^19,30^, which is too small to allow B_12_ passage in either conformation. This conformational change seems likely to contribute to the observed inhibition of locking by ATP analog binding to BtuCD by raising the energy barrier for the dual-lobe locking reaction that drives the complex into an asymmetrical conformation^31^.

We present evidence that the BtuCD-F locking reaction has a high activation enthalpy of ∼103 kJ/mol (Figure 3b-c), likely reflecting a substantial difference in protein conformation in the transition state for this reaction compared to the pre-reaction complex. The strong inhibition of this reaction by AMPPNP binding to BtuCD on the opposite side of the membrane from BtuF indicates that this conformational change involves coupled structural changes throughout BtuCD extending from the BtuF-binding interface in the periplasm to the ATPase active sites in the cytoplasm. We hypothesize that the global conformational transition mediating locking directly contributes to moving B_12_ through BtuCD during the transport process while simultaneous sealing the periplasmic surface of the transporter to prevent back-flow of B_12_ (Figure 6b). Passage of B_12_ requires the cavity at the interface between BtuC subunits to expand significantly compared to the crystal structures of BtuCD or the BtuCD-F complex determined in the absence of nucleotide or with AMPPNP bound to BtuD^19,31^ (**Extended Data Fig. 1** in ref. ^50^); this essential change in protein structure could occur in the high-energy conformation representing the transition state for the BtuCD-F locking reaction. Recent single-molecule, single-emission FRET-quenching studies of B_12_ transport BtuCD support this hypothesis^39^. A functional role for the locking reaction in B_12_ transport would rationalize the otherwise physiologically futile expenditure of energy when ATP is hydrolyzed in order to dissociate the locked BtuF-CD complex.

Given the evidence we have generated that ATP binding strongly inhibits locking (Figures 2d, 2e, & S3), the locked BtuCD-F complex is not likely to be found at a significant level in actively growing *E. coli* cells^44–47^ because they have a highly saturating millimolar-level concentration of ATP compared to the ∼8 µM *K*_*M*_ (Michaelis constant) of BtuCD for ATP hydrolysis^39^. Therefore, the BtuCD-F complex is likely to be found predominantly in the single-lobe anchored state in actively growing cells. Therefore, the anchored BtuCD•ATP-BtuF complex, represented by the BtuCD•AMPPNP-BtuF* complex characterized in this paper, likely represents the functional pre-translocation state of the transporter *in vivo*. Consistent with our hypothesis that the locking reaction plays a functional role the transport, the locked BtuCD-F complex is likely to represent the terminal stage of the translocation reaction. Additional studies, like those reported in a recent paper using single-molecule FRET studies to characterize the transport mechanism^39^, are needed to establish the role of the locking reaction in the transport of B_12_ by BtuCD-F.

## Acknowledgements

This work was supported by a RISE grant to JFH & RLG and graduate research fellowships to LZ, JK, and JER from Columbia University. LZ and CDK-T. were supported in part by the US National Institutes of Health Training Program in Molecular Biophysics (T32-GM008281), and CDK-T was also supported in part by the Graduate Fellowship Program of the US Department of Energy Office of Science (DEAC05-06OR23100). Additional support was provided by grants from the US NIH-NIGMS to JFH (GM127883) and RLG (GM084288 and GM137608) and grants from the US Cystic Fibrosis Foundation to JFH (HUNT18G0 and HUNT20G0). We thank D.C. Rees for provision of the WT BtuCD expression plasmid, Jia Ma and the Precision Biomolecular Characterization Facility at Columbia University for technical support, and the members of the Hunt and Gonzalez labs for scientific advice.

## Author contributions

LZ, JK, KL, JER, NKK, RLG, and JFH conceived the project and designed the experiments, which were performed by LZ, JK, KL, and JER and analyzed by LZ, JK, KL, JER, CKT, RLG, and JFH. LZ, JFH, and RLG drafted the manuscript and finalized it with help from all authors.

## Conflict of interest

JFH is a member of the Scientific Advisory Board of Nexomics Biosciences, Inc., and a consultant for Cyrus Biotechnology. The other authors declare no competing financial interests.

### Note

*The Supplementary Data contains 4 figures*.

## METHODS

### Plasmids and mutagenesis

A cysteine residue was appended onto the C-terminus of BtuF for fluorophore labeling via sulfhydryl-maleimide chemistry for both ensemble and smFRET experiments that measure fluorescent signal. It was followed by a hexahistidine (His_6_)-tag. Non-interacting BtuF (niBtuF) was created by mutating Glu-72 and Glu-202, crucial residues for BtuCD-F binding^54^, to alanine. For smFRET experiments employing surface-tethered BtuF*, in addition to the appended C-terminus cysteine and His_6_-tag, a (Gly_4_Ser)_2_ sequence was added and served as a spacer to allow maximum mobility of BtuF*, which was followed by an AviTag (Avidity LLC, Aurora, CO) at the end of the C-terminus for covalent biotinylation for surface immobilization. For two-color ensemble FRET experiment, a cystine-less BtuCD construct was created by replacing all native cysteine residues with serine for site-specific labeling using the plasmids listed below. Periplasmic residue Q111 was subsequently mutated to cysteine for fluorophore labeling. For all other experiments, WT BtuCD was used.

**Table 1.**
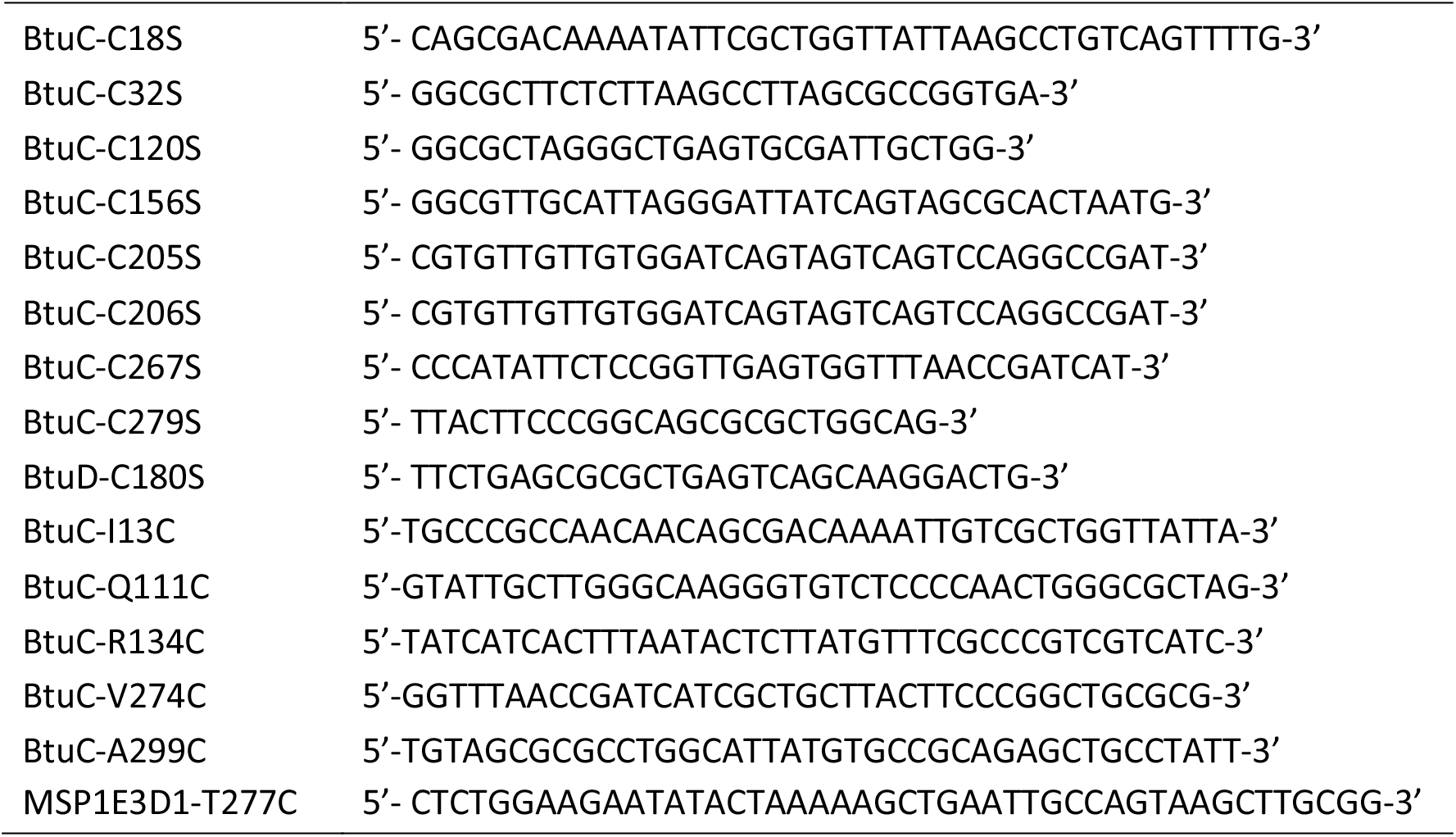
Primers used for mutagenesis.

### Protein expression, purification

Following induction at OD600=0.6 and overnight expression at 25 °C in *E. coli* C43 (DE3) cells (EMD Millipore, Novagen), cells were collected by centrifugation. All BtuF constructs were purified directly from whole cell lysate (obtained by sonication) via Ni-NTA chromatography followed by size-exclusion chromatography (Superdex 200 10/300 GL, GE Healthcare) in final buffers of 20-50 mM Tris-HCl, 100-150 mM NaCl, 0-1 mM Tris(2-carboxyethyl)phosphine (TCEP) at pH ∼7.4. All BtuCD constructs were purified from membrane pellets in 0.1% Lauryldimethylamine-oxide (LDAO) using the same purification techniques as above. Membrane scaffold protein (MSP) construct 1E3D1 that was used for nanodisc reconstitution was purified essentially as previously described^42^.

### Nanodisc and proteoliposome reconstitutions

Nanodisc-BtuCD was used in all experiments except for the liposome transport assay (Extended data Figure 2e-f). For nanodisc reconstitution, *E. coli* polar lipid extract (Avanti) was dried and rehydrated to 40 mg/mL (∼50 mM) in 100 mM NaCl, 1 mM TCEP, 100 mM sodium cholate and 20 mM Tris-HCl at pH 7.4 and mixed with MSP1E3D1^42^ and BtuCD. Final concentrations were: 5 mM lipid, 12 µM BtuCD, 72 µM MSP1E3D1, and 30mM sodium cholate. To minimize aggregation, 4% glycerol was also added. The mixture was incubated at 4 °C for 1 hour with gentle agitation. 400 mg of wet Bio-Beads SM-2 (BioRad) per 500 μL of mixture was added and the mixture continued to incubate overnight to remove detergents. Bio-beads were subsequently removed. Reconstituted nanodisc-BtuCD was purified from Ni-NTA agarose and then size-exclusion chromatography on a Superdex 200 column.

### Fluorophore labeling

BtuF was labeled with Alexa Fluor 546 (AF546) C_5_ Maleimide (ThermoFisher) according to the manufacturer’s manual. BtuCD Q111C was used in two-color FRET ensemble experiments and was labeled with Alexa Fluor 647 (AF647) C_5_ Maleimide prior to nanodisc reconstitution. All labeled proteins were purified from Ni-NTA agarose followed by size-exclusion chromatography to eliminate free fluorophores and aggregates.

### Ensemble FRET experiments in bulk solution

Fluorescence measurements in bulk solution (1.1 mL cuvette) were made on a fluorimeter (PC1 photon counting spectrofluorimeter; ISS), using an excitation wavelength of 540 nm and an emission wavelength of 573 nm to measure the intensity of AF546 on BtuF*. For single-color FRET experiments that use AF546-labeled BtuF and unlabeled BtuCD, total fluorescence was calculated from the formula *I*_*Total*_=*I*_*VV*_+2*G*\**I*_*VH*_ and anisotropy was calculated from the formula *r*=(*I*_*VV*_−*G*\**I*_*VH*_)/*I*_*Total*_, where *I*_*VV*_ is the fluorescence intensity recorded with excitation and emission polarization both in the vertical position, *I*_*VH*_ the fluorescence intensity with the emission polarization aligned in the horizontal position, and *G*=*I*_*HV*_/*I*_*HH*_. For two-color FRET experiments that used AF546-labeled BtuF and AF647-labeled BtuCD, additional fluorescent emission at 665nm was measured simultaneously.

### Kinetic analysis

Trapping of BtuF* by BtuCD in the presence of B_12_ was analyzed assuming the following reaction mechanism:

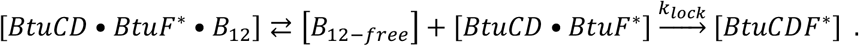

This model assumes that all BtuF* in the sample is associated with BtuCD in either a loosely associated or locked state, which is justified by the saturation of the change in the fluorescence anisotropy at the concentration of BtuCD used in the trapping experiments. Applying the steady-state assumption to the concentration of the reversibly associated BtuCD•BtuF* complex without B_12_ bound yields the following expression for the initial velocity of the trapping reaction in the presence of B_12_ :

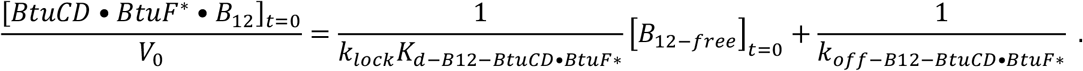

The variables *K*_*d−B12−BtuCD•Btuf*∗_ and *k*_*off−B12−BtuCD•BtuF*∗_ represent, respectively, the affinity constant for B_12_ binding to the reversibly associated BtuCD•BtuF* complex and the corresponding dissociation constant. The concentration of free B_12_ at time zero was calculated by subtracting from the total B_12_ concentration from the BtuF*•B_12_ complex concentration prior to the addition of BtuCD, which was calculated from the quenching level of BtuF*. The initial velocity was calculated from the slope of the kinetic progress curve in the first stable 15-sec window in each experiment, which was corrected for depletion of the initial complex based on the average BtuF* intensity in that region in order to yield the initial velocity.

## Supplementary data for

**Figure S1.**
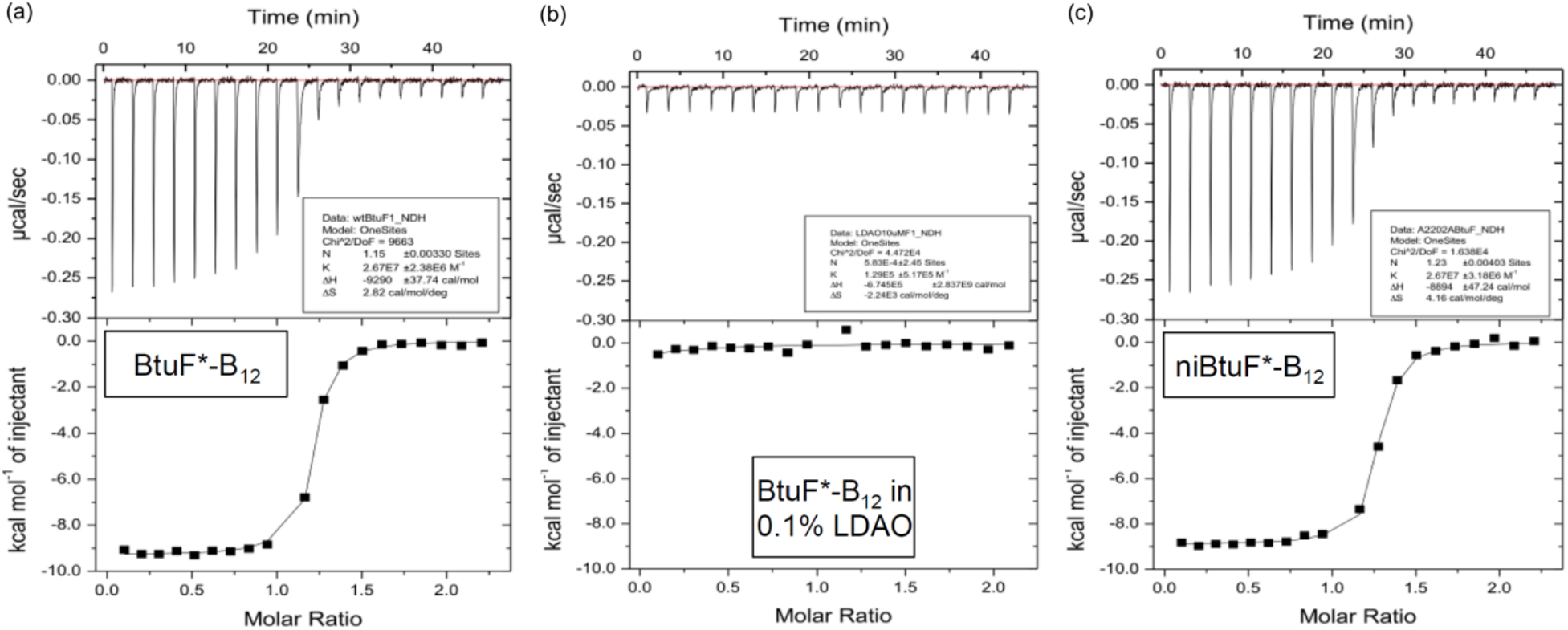
Isothermal calorimetry (ITC) measurements of B_12_ affinity for BtuF. **a-c**, Isothermal titration calorimetry measurements of B_12_ binding to BtuF*. The estimated values of *K*_*d*_ are 22 ± 10 nM for BtuF*•B_12_ and 54 ± 23 nM for non-interacting BtuF*•B12 (niBtuF*•B12). The niBtuF protein carries both E72A and E202A mutations, and it has no detectable affinity for BtuCD (data not shown). No binding of B_12_ to BtuF* is detected in the presence of 0/1% (w/v) LDAO). The uncertainty estimates give the standard error of the mean.

**Figure S2.**
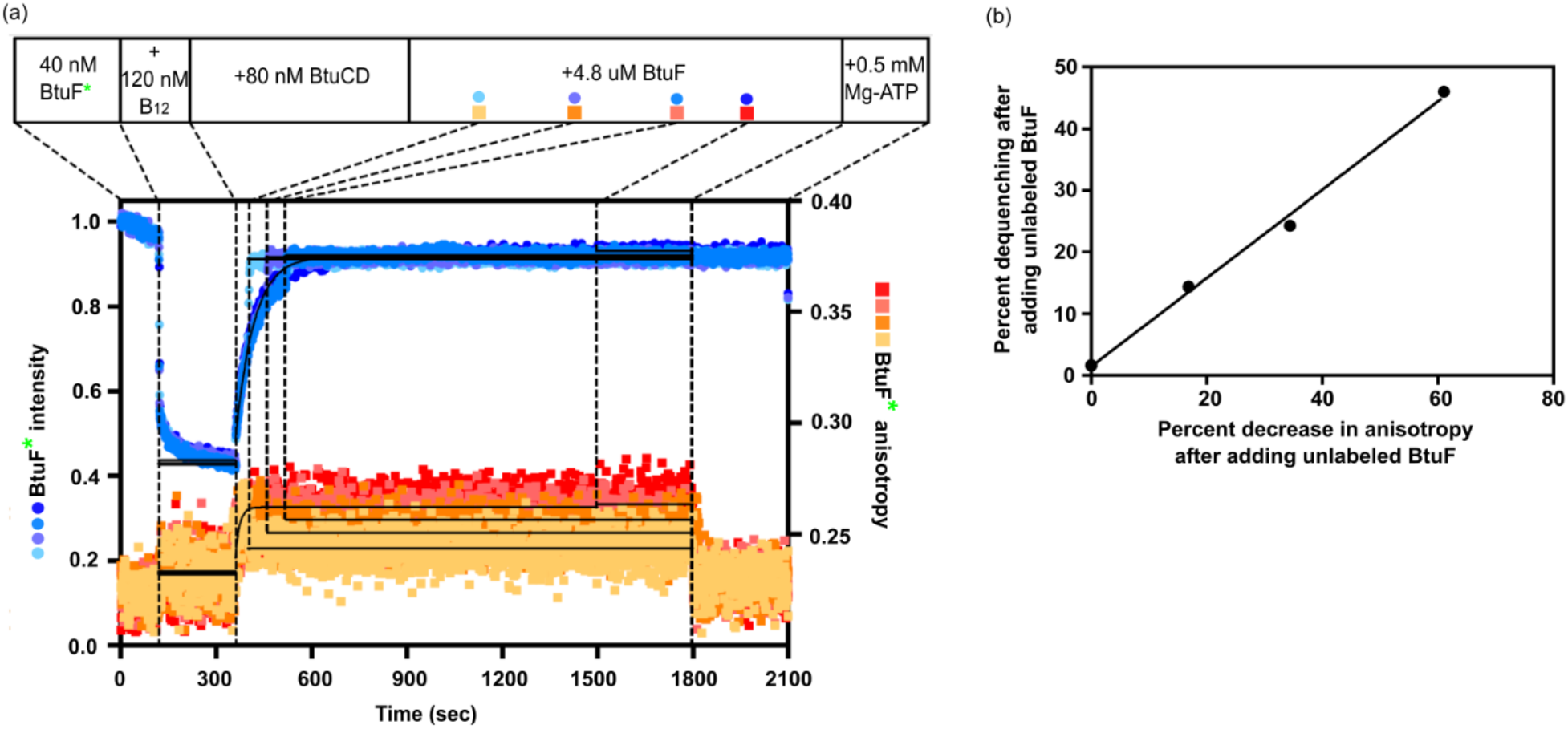
Further characterization of the BtuCD-F* locking reaction. **a**, Ensemble experiments measuring fluorescent intensity and anisotropy of BtuF* upon addition of B_12_, BtuCD, BtuF, and ATP at the beginning of the successive time windows. The different colors indicate different experiments performed with BtuF added at t = ∼400, 450, 500, or 1500 sec. **b**, The displaceable population of BtuF*, which is not locked onto BtuCD because B_12_ is bound, was quantified in the experiments in panel **a** by curve-fitting the decrease in anisotropy when excess unlabeled BtuF was added. The amount of BtuF*•B_12_ was quantified by the level of dequenching due to competitive binding of excess unlabeled BtuF to B_12_. The linearity of this plot indicates that formation of the non-displaceable or locked BtuCD-F* complex closely tracks release of B_12_ from BtuF*.

**Figure S3.**
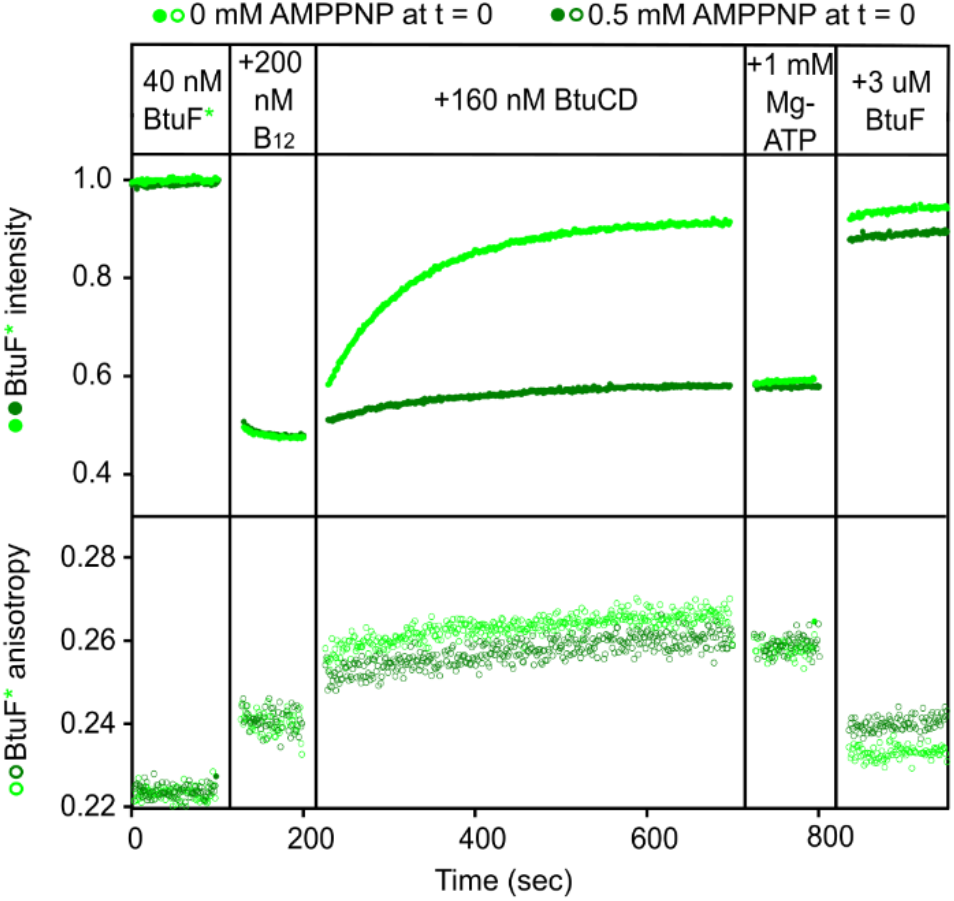
Further characterization of the influence of AMPPNP on BtuCD-F* locking. Ensemble experiments are shown that compare the gradual dequenching of BtuF* emission intensity upon addition of BtuCD to the BtuF*•B_12_ complex either in the presence (green) or in the absence of AMPPNP (dark green). This experiment was conducted at 25° C in the same buffer used in Figure 1 in the main text.

**Figure S4.**
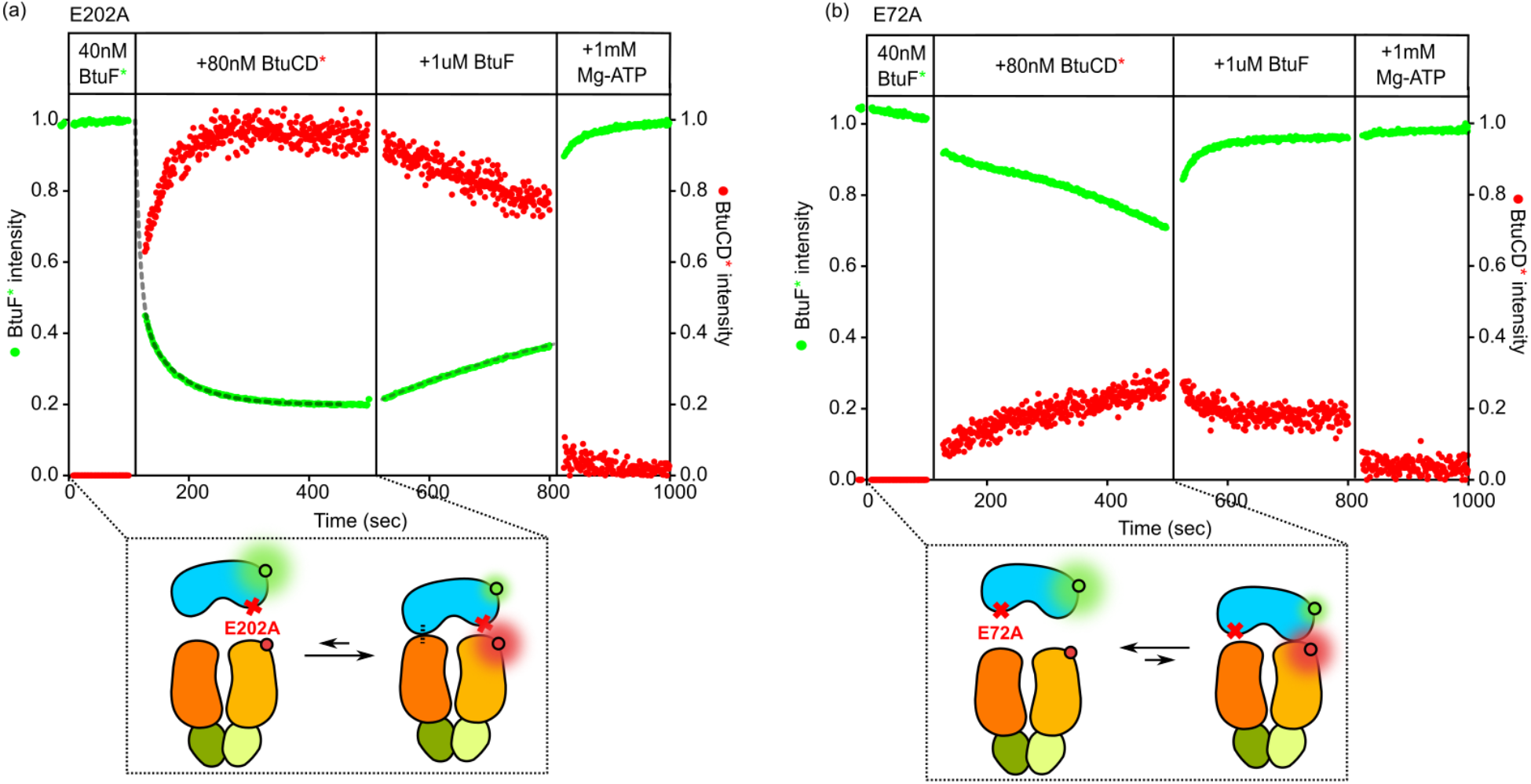
Two-color FRET experiments to characterize the roles of the two lobes of BtuF in mediating locking to BtuCD. **a**, An experiment is shown using an equivalent protocol to Figure 2d in the main text, but with E202A-BtuF* instead of WT-BtuF*. The change in green emission intensity upon the addition of BtuCD* was curve-fit using a double exponential function, yielding k_fast_ = 0.124 ± 0.002 sec^-1^ and k_slow_ = 0.016 ± 0.0001 sec^-1^ (corresponding to rate constants of 8.1 and 62 sec, respectively). **b**, An experiment is shown using an equivalent protocol to Figure 2d in the main text, but with E72A-BtuF* instead of WT-BtuF*. The experiments were conducted at 25° C in the same buffer used in Figure 1 in the main text.

